# Germinated brown rice research: bibliometric analysis of progress, hotspots and trends

**DOI:** 10.1101/2024.09.24.614820

**Authors:** Wenyang Zhang, Hao Niu, Yewang Zhao, Qiang Zhang, Fuxue Yang, Hong Zhang

**Author notes:** (W.Z.); (F.Y.). (Y.Z.). **Author Contributions:** Conceptualization, W.Z. and H.N; methodology, W.Z; software, W.Z; validation, H.N; formal analysis, Y.Z; investigation, Q.Z; resources, Q.Z; data curation, F.Y; writing original draft preparation, W.Z;writing review and editing,Y.Z. and Q.Z; visualization, F.Y. and H.Z; supervision, H.Z. and Q.Z; project administration, H.N; funding acquisition, H.Z. All authors have read and agreed to the published version of the manuscript. **Individual’s accomplishments:**Student from Tarim University,W.Z; Teacher from Tarim University,H.N; Teacher from Huazhong Agricultural University,Q.Z; Student from Tarim UniversityF.Y; Dean of College of Mechanical and Electronic Engineering,Tarim University,H.Z. **Ethics approval:** Not applicable. **Consent to participate:** All co-authors have agreed with the contents of the manuscript. **Consent for Publication:** All co authors agree to publish. **Availability of data and material:** All data generated or analysed during this study are included in this published article. **Funding:** Alaer Science and Technology Program “Development and Application of Key Technology and Equipment for Germinated Brown Rice Processing” (Grant No:2023ZB01). All authors have seen the manuscript and approved to submit to your journal. This paper is new. Neither the entire paper nor any part of its content has been published or has been accepted elsewhere.

## Abstract

To gain a deeper understanding of global research trends and focal points in germinated brown rice, this article takes the relevant literature on germinated brown rice in the core database of Web of Science as the research object. By using bibliometric analysis, the literature on germinated brown rice published from 2003 to 2023 is deeply analyzed, and the global research progress, hotspots and future development trend of germinated brown rice are summarized. Since 2018, research on germinated brown rice has been rapidly developing, with a significant surge in interest since 2020. China leads in the number of publications, institutions, and core authors in this research area. The primary research topics include the nutritional value, physiological active components, and industrial applications of germinated brown rice. Current research frontiers involve identifying, evaluating, and enhancing bioactive substances in germinated brown rice for food applications. Research in this field remains active, and application scenarios are becoming increasingly diverse. Future studies may explore new directions in related equipment and ingredients.

## Introduction

Germinated brown rice is produced through a series of processes including hulling, impurity removal, screening, grading, washing, soaking, germination, drying, and testing (Zhang and Liang. 2019). Current research on the production technology of germinated brown rice primarily focuses on the enrichment, soaking, and drying of γ-aminobutyric acid (GABA), with GABA enrichment being the most studied aspect (Yang et al. 2017). GABA enrichment process is the focus of germinated brown rice research, and its detection method is the basis of all research work (Chen et al. 2023; Wang et al. 2016). GABA can regulate emotions, improve sleep quality and enhance the brain activity (Liu et al. 2023). At the same time, its content in germinated brown rice is much higher than that of brown rice (Zhang. 2014). The humidification process of germinated brown rice is directly related to its quality, with more even mixing leading to better humidification uniformity (Cao et al. 2015). The drying process also significantly affects the content of main nutrients, enzymatic activity, and hardness of germinated brown rice (Sun et al. 2022).

The bibliometrics used in this paper are based on the research of literature system and metrology characteristics, and uses mathematical and statistical research methods to deeply explore the quantity, distribution and change of literature, and thereby reveal the internal structure, characteristics and laws of science and technology (Fan et al. 2009). Since the beginning of the 20th century, the quantitative research of literature has begun to sprout. After nearly a century of development, bibliometrics has gradually matured, and its theory and practical application have been extensively and deeply studied. During this period, many outstanding scholars and works have emerged continuously, making important contributions to the development of bibliometrics. However, with the rapid development of the Internet in today’s society, the research methods of bibliometrics have undergone significant changes, and its research hotspots have also changed (Zhao et al. 2010).

This paper uses bibliometrics to analyze the literature on germinated brown rice research in the core database of Web of Science from 2003 to 2023, and analyzes the current situation, hotspots and future trends of germinated brown rice research from a global perspective. By reviewing and visualizing the development process of germinated brown rice research on a time scale, this paper aims at provide researchers with a deep understanding of the development characteristics of this field. It is hoped that this will provide more valuable reference and guidance for future related research (Xiao et al. 2020; Lan et al. 2022).

## Material and methods

### Material

This article is indexed in the Web of Science core collection using the search formula TS=(“brown rice” OR “germinated brown rice”) AND TS=(“process” OR “enzyme” OR ‘‘γ-aminobutyric acid’’ OR ‘‘humidification’’ OR ‘‘machine”) NOT TS=(‘‘brown planthopper’’ OR ‘‘disease”). The language was set to English, the literature published from January 1, 2003, to October 1, 2023, was searched. A total of 1441 search results were obtained. The search results will be slightly changed due to changes in the database, and the change in the number of documents has little impact on the final conclusion of the paper. The data search ended on November 1, 2023.

## Methods

The data processing software Origin (2021) and bibliometric analysis tool CiteSpace (6.1.R6), Tableau (Public 2021.3), Charticulator (https://charticulator.com/app/index.html), and VOSviewer (1.6.20) to retrieve the data for quantitative analysis. Origin, Tableau and Charticulator software were used to analyze the number and trends of germinated brown rice studies. Knowledge graph visualization software CiteSpace and VOSviewer were used to analyze the institutions and authors with high frequency of publication volume, high frequency keywords, keyword co-occurrence map, breakout map and time series map (Lan et al. 2022). The temporal and spatial development of germinated brown rice research was reviewed and visualized.

### General overview of research related to germinated brown rice

#### Study time distribution

Figure 1 illustrates the quantity and timing of publications in the top 10 countries or regions in the field of germinated brown rice research. The upward trend in publication numbers over time suggests that this area of study is gaining global attention. However, due to the time constraints of the search, the number of published papers in 2023 was incomplete, so the number of published papers in this year decreased to a certain extent.

**Fig 1.**
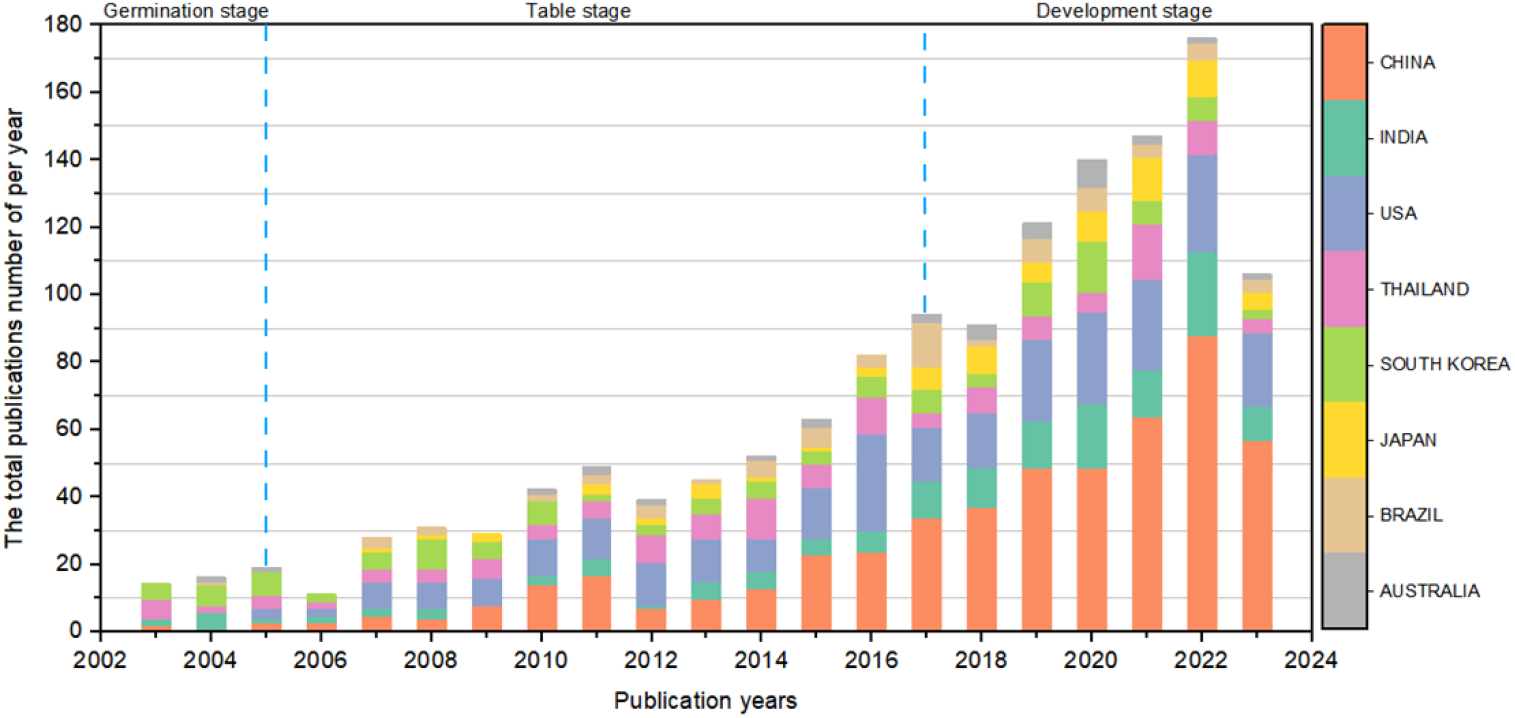
Cumulative histogram of annual changes in the number of publications in the top 10 countries or regions in the field of germinated brown rice research

Research trends indicate a sharp increase in global studies on germinated brown rice since 2017, making it an increasingly focal point of research. These data can be primarily divided into three stages: germination stage (2003-2005), table stage (2006-2017) and development stage (2018-2023).

(1) In 2003, only 16 papers were published. The number of publications increased to 20 in 2004, but has since declined.
(4) Between 2006 and 2017, the overall number of papers published was in a sustained growth trend, but the growth trend was tortuous. The number of published papers peaked at 94 in 2017, after which it declined.
(3) Since 2018, the proportion of germinated brown rice research has increased, mainly reflected in the number of papers published in the field of germinated brown rice. From 2018 to December 1, 2023, approximately 750 papers were published, with this section research accounting for about 53% of the total. This indicates that the research on germinated brown rice began to enter a stage of rapid development in 2018 and has been growing rapidly since then.This trend suggests that the research in this field is expected to continue increasing in the future, demonstrating sustainable development trend.

In summary, research on germinated brown rice has experienced rapid growth in the past few years, and the number of studies is expected to continue increasing in the future.

#### Study spatial distribution

##### Country or region

A country’s scientific research strength and academic influence can be reflected to some extent by its total output of papers. Moreover, a paper’s citation frequency serves as a crucial metric for gauging its global reach and significance within the discipline (Xiao et al. 2020). According to the core database of Web of Science, the distribution of papers published between 2003 and 2023 is illustrated in Figure 2(a).

**Fig 2.**
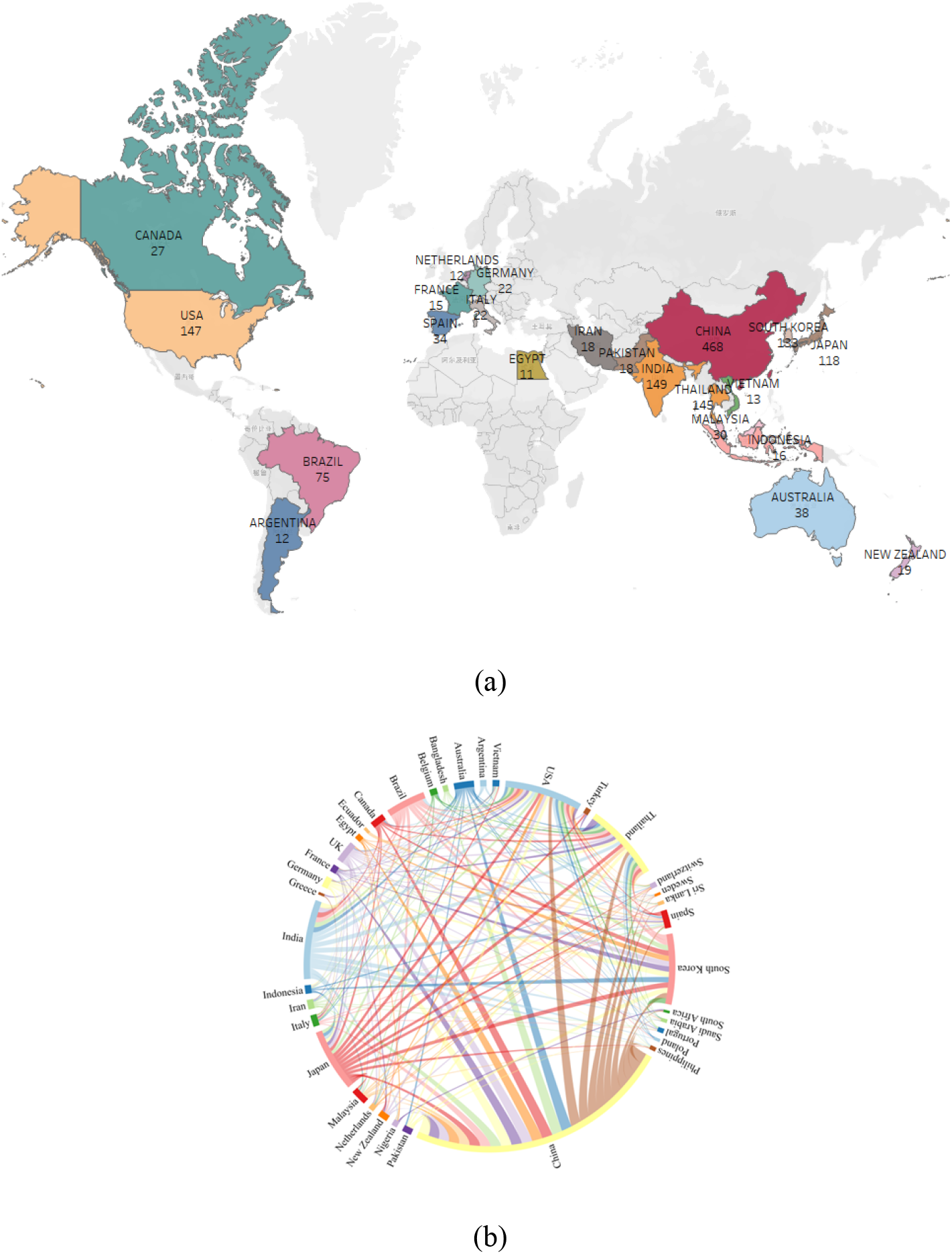
**(a)** Main source country or region of literature and its distribution map; **(b)** Cooperation between countries or regions

China, India and the United States are currently the leading countries in germinated brown rice research. China has published 468 papers, accounting for approximately 32.98% of the total; India has published 149 papers, accounting for about 10.50%; and the United States had published 147 papers, accounting for about 10.36%. It is also known that Japan and South Korea were the earliest countries to study germinated brown rice. Suzuki, K (2003) developed a new technique for germinating brown rice by using a twin-screw extruder to extrude germinated brown rice, which contains higher GABA compared to white and brown rice (Suzuki et al. 2003). Additionally, South Korea, Thailand and Japan are also countries where germinated brown rice research is more active. To better understand the research status of germinated brown rice in different geographical regions, all countries or regions were divided according to geographical regions, and the spatial distribution of literature related to germinated brown rice research was obtained. Asia and North America are the main research areas for germinated brown rice; within these regions, the prominent countries include the agricultural powerhouse China, the populous India, and the agriculturally advanced United States.

To deepen understanding of country collaboration in the field of germinated brown rice research, the Charticulator tool is used to conduct a systematic analysis of published country and regional cooperation and present the results in Figure 2(b).

According to the analysis of co-occurrence relationships among the top countries or regions in the number of published articles, China, Germany, the United States, the United Kingdom, Canada, and Australia are the most closely connected countries, indicating that these countries are the most frequent cooperative relationships in the field of germinated brown rice. It can be seen that China has carried out more cooperation with the main publishing countries or regions. This may be attributed to China’s sustained and firm investment in agriculture, rural areas and farmers, and the growth rate of fixed asset investment in the primary industry is significantly higher than the average growth rate of fixed asset investment in the whole society (Wang. 2019). Chinese scholars are increasingly securing valuable opportunities for deep exchanges and collaborations with top global institutions and scholars. Such interactions will undoubtedly propel domestic germinated brown rice research to the international forefront and hotspots, fostering collective advancement in scientific research.

In conclusion, Asia holds significant influence in germinated brown rice research, particularly in publications, with North America, represented by the United States, also making a notable impact. Moreover, research related to germinated brown rice in other countries and regions is demonstrating an upward trend.

##### Institutions or units

According to the data in the core collection of Web of Science, Table 1 shows the top 15 institutions or units in the field of germinated brown rice.

**Table 1.** Institutions or units that produce journal papers.

| Rank | Units/institutions | Countries | Publications | Ratio/% |
| --- | --- | --- | --- | --- |
| 1 | Ministry of Agriculture and Rural Affairs of China | China | 49 | 3.453 |
| 2 | Karlstad University | Sweden | 42 | 2.960 |
| 3 | Chinese Academy of Sciences | China | 40 | 2.819 |
| 4 | Jiangnan University | China | 40 | 2.819 |
| 5 | Chinese Academy of Agricultural Sciences | China | 39 | 2.748 |
| 6 | Zhejiang University | China | 33 | 2.326 |
| 7 | United States Department of Agriculture | USA | 29 | 2.044 |
| 8 | King's University of Science and Technology,<br>Thailand | Thailand | 26 | 1.832 |
| 9 | Nanjing Agricultural University | China | 26 | 1.832 |
| 10 | China Agricultural University | China | 25 | 1.762 |
| 11 | Chiang Mai University | Thailand | 23 | 1.621 |
| 12 | Huazhong Agricultural University | China | 23 | 1.621 |
| 13 | Japan National Agri-Food Research Organization | Japan | 21 | 1.480 |
| 14 | Pelletas Federal University | Brazil | 21 | 1.480 |
| 15 | Yangzhou University | China | 21 | 1.480 |

There are four institutions that have published 40 or more papers, with two from China and one from Sweden. The Ministry of Agriculture and Rural Affairs of China has published the most papers, totaling 49, which accounts for 3.453% of all publications. This is followed by Karlstad University with 42 papers, and both the Chinese Academy of Sciences and Jiangnan University with 40 papers each.

China dominated the top 10 list, securing 7 positions. The Ministry of Agriculture and Rural Affairs of China, the Chinese Academy of Sciences, and Jiangnan University emerged as the leading institutions in germinated brown rice research. The Chinese Ministry of Agriculture and Rural Affairs is mainly concerned with food science and technology and environmental science ecology, such as the review of the latest literature on the physiological functions, preparation methods, enrichment methods, metabolic pathways and applications of GABA (Zhou et al. 2022), while the Chinese Academy of Sciences is mainly concerned with environmental science ecology and plant science. For example, rapid viscosity analysis, differential scanning calorimetry and X-ray diffraction were used to analyze the structural, physicochemical and functional changes in the germination process of brown rice (He. 2022), while Jiangnan University mainly focused on food science and technology and chemistry. For example, brown rice of three different rice varieties was germinated by aeration and calcium ion germination treatment. The research results show that germination treatment is an effective method to produce brown rice with appropriate edible quality and function (Zhang. 2022).

Figure 4 illustrates the collaborative relationships among the aforementioned agencies or units. Universities in China, including Zhejiang University, China Agricultural University, Nanjing Agricultural University, and Jiangnan University, frequently engage in cooperative endeavors. However, collaborations with foreign institutions or units are less common. Notably, Kasetsart University spearheads significant cooperation efforts. To further advance this field, augmenting interactions with diverse national institutions, educational establishments, and corporations is imperative, as is reinforcing international partnerships in the domain of germinated brown rice.

#### Research progress and hotspots of germinated brown rice

Keywords in academic papers are labels that indicate the core of the research and subject information. They provide a higher level summary of the main content of the paper, which can be used to identify the main content of the paper and the academic research direction, and analyze the keywords of germinated brown rice research in different directions (Fan. 2009). On this basis, the whole research progress in the field of germinated brown rice is analyzed, the current research hotspots are identified, and the future development trend is predicted.

#### Keywords co-occurrence analysis

According to the literature, the research of germinated brown rice covers many aspects. Through co-occurrence analysis of high-frequency keywords in the literature, this section identifies relevant key contents and research questions. Based on the co-occurrence analysis of keywords, a color-differentiated co-occurrence graph is generated to investigate the contribution of keywords in the literature and the close relationship between keywords, as shown in Figure 4. In the co-occurrence graph, centrality refers to the magnitude of the linking role of nodes in the network, which reflects the influence of the co-occurrence relationship in the co-occurrence graph. The stronger centrality indicates that researchers pay more attention to such keywords, and the content closer to the co-occurrence word center is more important.

The keywords with the highest frequency occurrence according to co-occurrence graph analysis included brown rice, rice, quality, physicochemical properties, and antioxidant activity. Brown rice and rice appeared the most frequently (449 times), indicating that germinated brown rice and brown rice were the main background of the study. The former mainly investigated the effects of amendments on cadmium accumulation in germinated brown rice, GABA content in germinated brown rice cooked at different times, antioxidant capacity, amino acids produced by 15 proteins, TP and reducing sugar content and color change, and slightly acidic electrolytic water treatment on GABA accumulation in germinated brown rice (Li et al. 2022; Tomoyasu et al. 2021; Li et al. 2015). The latter studies mainly focused on the combined process of GABA content in low-pressure steam brown rice and low-pressure superheated steam dried brown rice, changes in physical and enzymatic digestion behaviors of cooked white rice and brown rice with similar amylose content, nutritional quality of whole grain brown rice, emphasizing the main bioactive components of brown rice (Yan et al. 2022; Wu et al. 2017; Mohammed and Bin. 2023).

#### Keywords cluster analysis

Through the cluster analysis of the keywords co-occurrence network, the log-likelihood ratio (LLR) algorithm is used to extract the cluster labels, and a total of 10 cluster labels are obtained. The smaller the cluster sequence number, the more keywords the cluster contains. That is, #0 is the largest cluster, and so on.The clustering result analysis is shown in Figure 5.

**Fig 3.**
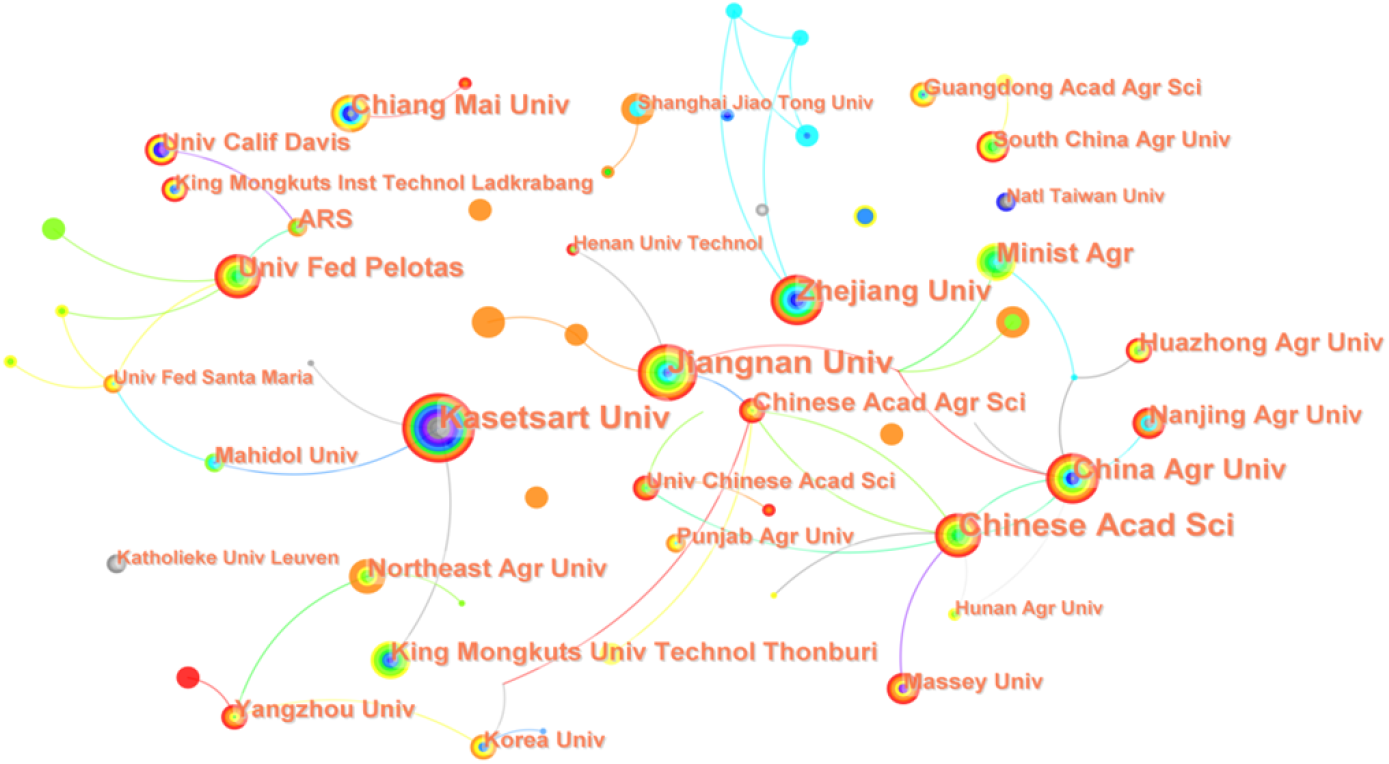
Agency or unit collaboration diagram

**Fig 4.**
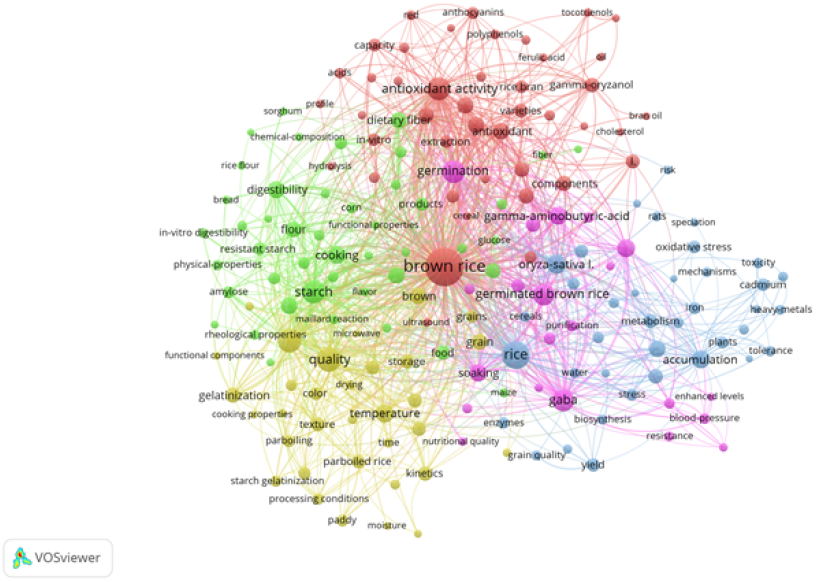
Keywords co-occurrence graph

**Fig 5.**
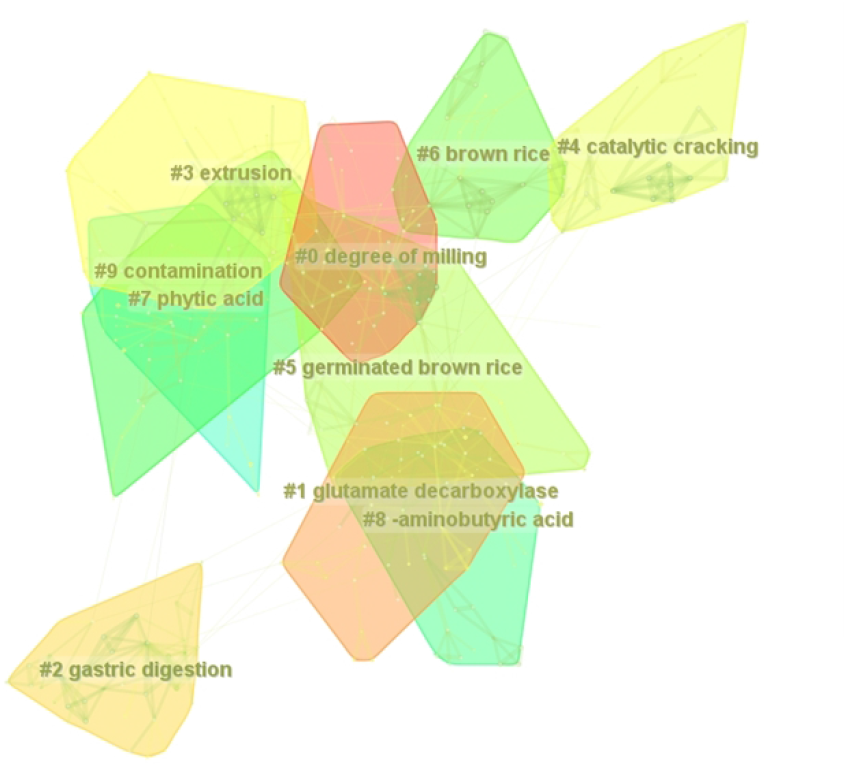
Keywords clustering diagram

Based on the results of cluster analysis and the development context of germinated brown rice, the distribution of 10 cluster words was summarized into 3 themes. The research contents include #1 glutamate decarboxylase, #5 germinated brown rice, #6 brown rice, #7 phytic acid and #8 gamma-aminobutyric acid; Research methods include #3 extrusion and #4 catalytic cracking; The research indexes are #0 degree of milling; The types of research include #2 gastric digestion and #9 contamination. It can be seen that the research on germinated brown rice mainly focuses on the nutrition composition and processing technology.

The keywords clustering time graph shows which keywords are included in each cluster, and the time nodes of the beginning and end of the clustering topic, reflecting the distribution time span and importance of a certain cluster (Liu et al. 2022), thus summarizing the evolutionary path of germinated brown rice. The time line graph pays more attention to the relationship between keywords clusters (Fig. 7). Cluster topics #8 gamma-aminobutyric acid and #7 phytic acid are associated with a large number of keywords, indicating that GABA and phytic acid are hot directions for nutrient composition related research of germinated brown rice. The clustering theme #2 gastric digestion has a long gastric digestion time span, which indicates that health-related topics such as stomach digestion and digestive system are highly and persistently popular in the development and research of germinated brown rice.

#### Keywords breakout analysis

In this paper, CiteSpace is used to carry out emergent analysis of keywords in related research. Emergent words refer to the words whose value of keywords rises sharply and becomes a hot topic in a short period of time (Liu et al. 2022). The intensity of a keywords’ emergence is directly proportional to the activity of the topic it represents in the corresponding time period. Red colour means high heat, blue colour means low heat, and the higher the value of strength means the more important the keyword. The Top 15 emergent keywords are shown in Figure 6. These emergent keywords are identified as those with wide interest and great influence in the academic field.

**Fig 6.**
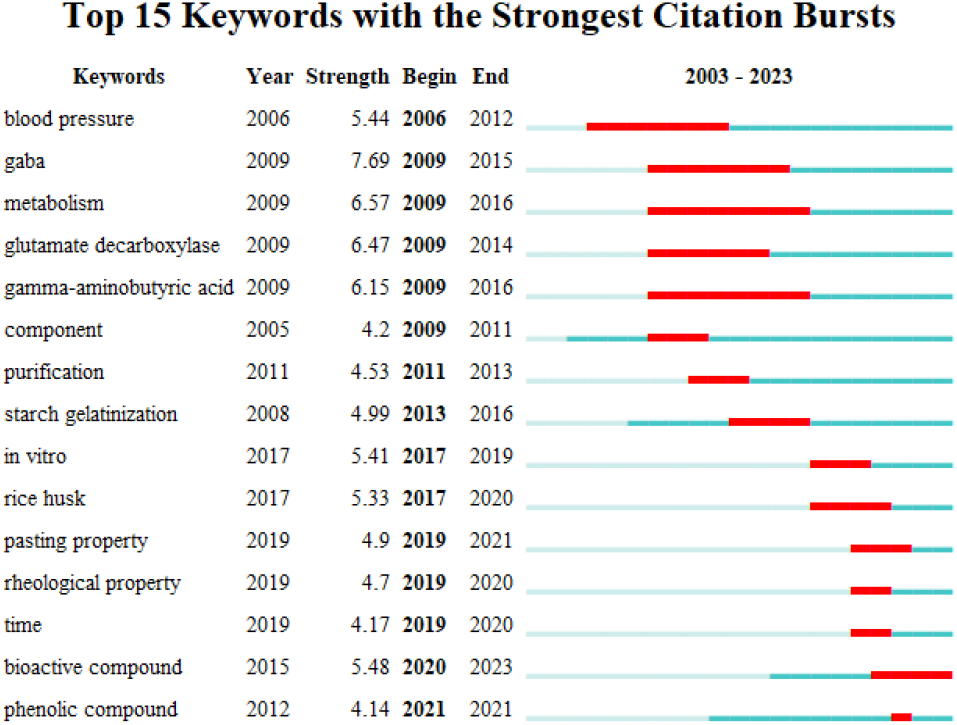
The evolution of hotspots of emerging keywords

**Fig 7.**
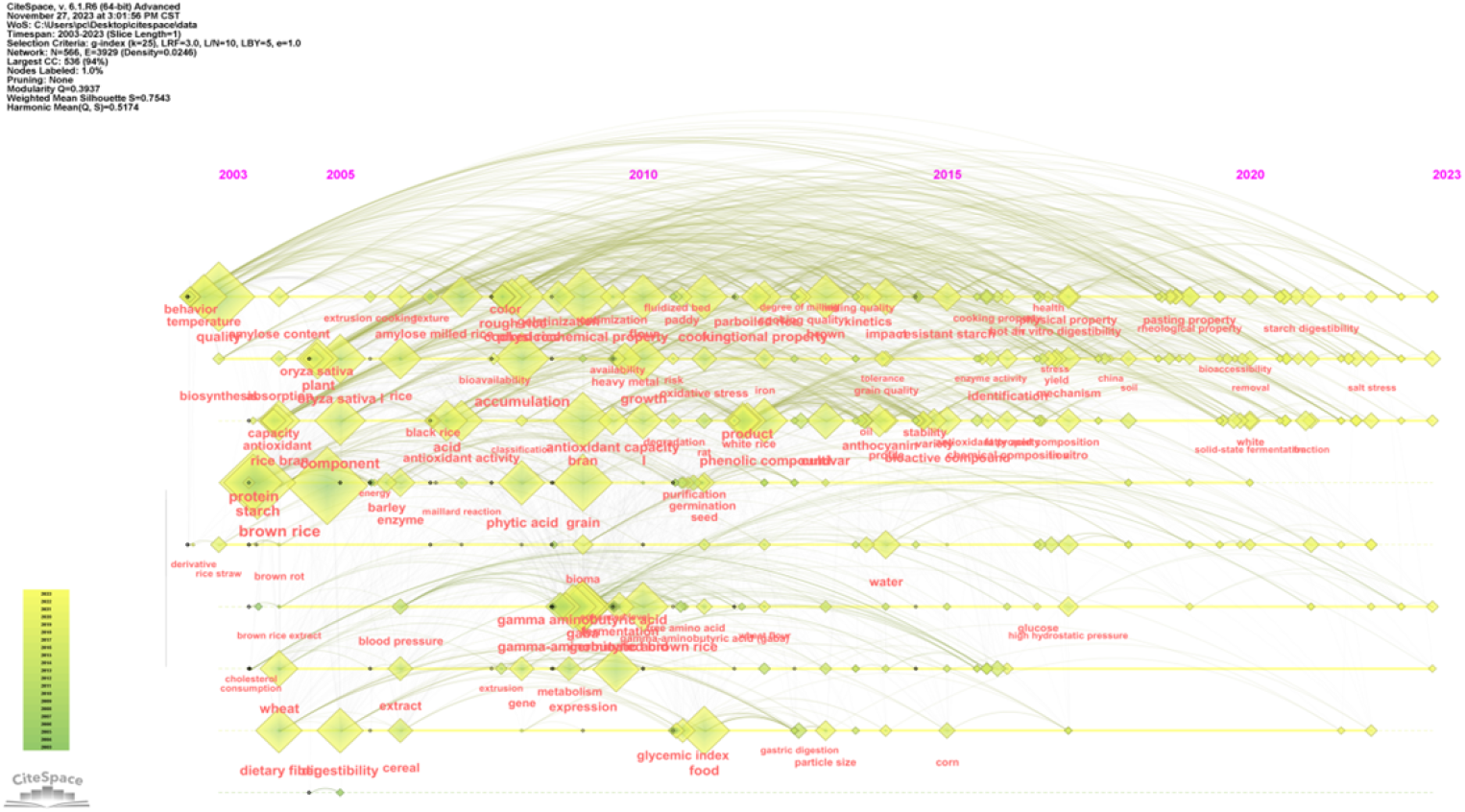
Time sequence diagram of keywords

With the change of time, the research object gradually expands and the research content becomes more diversified. From 2006 to 2016, the research mainly focused on human health aspects such as blood pressure (Odahara et al. 2008)metabolism (Chung et al. 2009), and gradually began to explore GABA and starch gelation (Wang et al. 2020). During 2017-2023, its internal components such as rheological properties (Zhu et al. 2019), bioactive compounds (Ohtsubo et al. 2005; Baek et al. 2005; Moongngarm and Saetung. 2010)and phenolic compounds and other sudden words are highly popular, indicating that the research focus has gradually shifted to the study of the internal properties of germinated brown rice, which may be due to the continuous optimization of relevant detection equipment, and then promote the study of internal nutrients to be more in-depth. Therefore, the research should pay more attention to its internal components, in order to further study its food safety and health effects.

#### Keywords timing analysis

The LLR algorithm in CiteSpace software was used to visualize the time line analysis of keywords in the literature to further clearly and intuitively investigate the development history of germinated brown rice research. The time series diagram describes the keywords contained in the cluster in terms of time. The density of the lines among the keywords represents the frequency of occurrence, and the lines between the keywords represent the connections between them. Finally, the development context of the field of germinated brown rice is obtained. Combined with the number of papers published and the occurrence of keywords, the research vein and guiding point are further obtained according to the time node and keywords. The keywords in Figure 7. are the years in which they first appeared in the data set.

The keywords co-occurrence chart is divided into three stages according to the time axis: early stage (2003-2005), middle stage (2006-2017) and 2018-2023, and compares and analyzes the research context and frontier of germinated brown rice.

In the early stage, the research on germinated brown rice began based on brown rice, and was located in the fields of its behavior temperature, quality, protein starch and amylose. Similar to degradation of soluble protein of brown rice by endogenous proteolytic activity (Yamada et al. 2014), high-precision sensing of brown rice freshness over time by chemiluminescence measurement and luminol peroxidase system (Noda et al. 2005), construction of single grain brown rice quality sorting system based on near infrared (Rittiron et al. 2004); In the middle period of the study, the keyword GABA appeared for the first time, and GABA has been the focus of attention in subsequent studies, such as the influence of slightly acidic electrolytic water, soaking and gas treatment on GABA content (Li et al. 2015; Komatsuzaki et al. 2007) and the self-reinforcing effect of GABA in rice bran under different stress conditions (Kim et al. 2015). At the same time, various production processes and evaluation indicators of germinated brown rice are more perfect and diversified, such as the study of the optimal water addition rate of brown rice during the germinating period by segmented humidity regulation (Cao et al. 2015). The optimization of millimeter wave and cellulase processing parameters (Seo et al. 2016; Zhang,Q et al. 2015); from 2018 to 2023, the related keywords are gradually reduced, and the main research contents are membrane reactor (Sitanggang et al. 2021), high temperature cooking (Toyoizumi et al. 2021), pulsed light (Zhang et al. 2022), static magnetic field treatment (Luo et al. 2023), and magnetic field treatment (Chen et al. 2023). This stage is basically the update and improvement of the previous research content and process, and there are not many hot topics of development technology.

The current research on germinated brown rice primarily concentrates on the advancement of processing technology and nutritional components. However, there is a growing emphasis on understanding the raw quality and its influence on human health. Despite this, interest in germinated brown rice has waned in recent years, presenting several challenges. Firstly, the specific alterations in GABA content remain unclear, and there is a paucity of research on the drying process. Additionally, the optimal integration of humidification and conditioning processes with GABA enrichment to produce high-GABA germinated brown rice requires further investigation. Secondly, there is limited research on specialized processing equipment that aligns with these procedures, leading to difficulties in controlling product quality, complex processes, poor equipment integration, extended research and development cycles, and elevated costs.

## Discussion

This study identified global journal papers on germinated brown rice published in the Web of Science database between 2003 and 2023. The quantitative distribution and sources of relevant studies were analyzed using bibliometrics. Based on the analysis atlas, the research content, research progress, main hotspots and development trend of germinated brown rice were summarized. The main conclusions of the content are summarized as follows:

(1) From 2003 to 2005, the research on germinated brown rice was nascent, with a limited number of countries involved and consequently, few published papers. Over time, there was a gradual increase in both the number of relevant papers and the countries participating in the research. Between 2006 and 2017, the field experienced steady growth as research papers continued to be published. From 2018 to 2023, there was a rapid surge in the number of studies. It is anticipated that the publication count in this field will continue to rise in the subsequent years.
(2) Research on germinated brown rice predominantly takes place in Asia, the Americas, and Europe, with China being the most active participant. Furthermore, there is significant collaboration between China, Thailand, Japan, and the United States, as well as between South Korea and Japan. Such international cooperation fosters the diverse development of germinated brown rice research. The support from these nations not only aids research institutions and teams in acquiring essential resources but also motivates researchers to delve deeply into various aspects of the field. This multifaceted research approach is likely to yield numerous scientific advancements and technological innovations that will cater to the demand for germinated brown rice.
(3) The following research institutions are the major research force to papers in the field of germinated brown rice: Chinese Ministry of Agriculture and Rural Affairs, Chinese Academy of Sciences, Jiangnan University, and Karlstads University. Among them, Ministry of Agriculture and Rural Affairs of China has a significant contribution and influence in this field.
(4) In the reviewed literature, GABA is frequently used as a primary evaluation metric. The predominant processing techniques encompass humidification, soaking, drying, exposure to magnetic fields, and microwave treatment. In recent years, research into the nutritional components and processing methodologies of germinated brown rice has garnered increased attention, marking a new phase in its development.

Furthermore, as modern technologies such as big data and artificial intelligence continue to evolve, these advancements are expected to foster the maturation of research techniques pertaining to germinated brown rice. Simultaneously, there is potential for these technologies to be integrated into the entirety of the production and processing stages for various grains, aiming to fulfill market demands for grain-based products.

Utilizing official internet data, this study compiles recorded data on germinated brown rice spanning from 2003 to 2023. Nonetheless, it is crucial to acknowledge that incorporating multiple databases may introduce issues such as sample overlap, despite the potential for some disciplines to be concurrently included in other databases, which could result in data omissions.

## Abbreviations

GABA Γ-: aminobutyric acid
NIRS: Near infrared spectroscopy
WD: Wavelet denoising
IF: Impact factor
JCR: Journal citation reports
LLR: Log-likelihood ratio

